# Allelic Variation at 9p21.3 Orchestrates Widespread RNA Splicing Shifts Governing Vascular Smooth Muscle Cell Plasticity

**DOI:** 10.64898/2026.01.14.699552

**Authors:** S. Suryavanshi, H. Yang, E. Salido, V Lo Sardo

## Abstract

Genetic risk for coronary artery disease (CAD) has been linked to variants across more than 300 genomic loci. Whether and how these loci interface with RNA processing to drive disease-relevant cellular phenotypes remains unknown. Here, we applied haplotype-biased genome editing in induced pluripotent stem cells (iPSCs), followed by differentiation into vascular smooth muscle cells (VSMCs), to address how genetic variation at the strongest CAD locus – the 9p21.3 CAD risk locus – affects RNA processing and alternative splicing genome-wide. Using long-read RNA sequencing, we identified distinct allele-specific transcriptional programs driven by the two major haplotypes at 9p21.3, risk and non-risk. We unravel extensive reprogramming of mRNA splicing across the transcriptome, which leads to VSMC aberrant phenotypic modulation. The 9p21.3 risk haplotype disrupts transcript isoform expression and usage across multiple genomic loci implicated in diverse stages of atherosclerotic plaque development. We prioritized DDX5, previously implicated in CAD through GWAS. Isoform-specific modulation of DDX5 in VSMCs was sufficient to mitigate the 9p21.3 risk-associated molecular signature. Together, this work provides the first comprehensive isoform-level transcriptomic comparison of the two major haplotypes at the 9p21.3 locus and identifies a 9p21.3–DDX5 axis as a key regulator of VSMC phenotypic plasticity. These findings uncover allele-specific reprogramming of RNA splicing as a previously unrecognized mechanism underlying cardiovascular disease susceptibility and present a resource of targetable transcripts with potential relevance across vascular pathologies.

## INTRODUCTION

Coronary artery disease (CAD) remains a leading cause of morbidity and mortality worldwide, with a significant genetic contribution to disease risk^1–6^. Genome-wide association studies (GWAS) have identified more than 300 loci associated with CAD susceptibility, many of which implicate vascular cell types central to atherogenesis, including vascular smooth muscle cells (VSMCs)^7,8^. Despite this progress, translating CAD-associated genetic variation into mechanistic insight remains a major challenge, particularly for non-coding loci whose effects are subtle, context-dependent, and difficult to model experimentally.

A growing body of evidence suggests that alternative RNA splicing and transcript isoform regulation play central roles in cardiovascular development and disease, including atherosclerosis^9–11^, vascular remodeling^12^, and heart failure^13,14^. Perturbations in splicing regulators such as RBFOX1^13^, RBM20^14^, SRSF3^15^, and CELF1^16^ is known to impair cardiac or vascular function. Nevertheless, how CAD-associated genetic loci—particularly non-coding loci—interface with RNA processing to shape vascular cell phenotypes remains poorly understood.

Here, we address this gap by combining allele-biased editing at the strongest CAD-associated locus, the 9p21.3, isogenic VSMC models carrying different alleles, and long-read RNA sequencing to interrogate how non-coding genetic risk factors shape cellular state in CAD.

The 9p21.3 locus — a 60 kb region on human chromosome 9 — stands out as the most robust and reproducible genetic association identified to date ^17–21^. This primate-specific locus contains ∼80 tightly linked variants that segregate into two major haplotypes conferring increased or reduced CAD risk across populations^17–19,21,22^. The risk haplotype accounts for an estimated 10-15% of CAD incidence in the United States^23^, underscoring its outsized clinical impact. Yet, despite nearly two decades of investigation, the molecular mechanism by which 9p21.3 confers disease risk remain incompletely understood. A key barrier has been the absence of an appropriate in vivo model, as the syntenic region in mice lacks the human haploblock structure and a clearly conserved homolog of the long non-coding RNA ANRIL encoded within the locus^24,25^.

To overcome these limitations, we have leveraged human induced pluripotent stem cells (iPSCs)-based systems to interrogate the 9p21.3 function in disease-relevant vascular cell types^26^. Using isogenic iPSC-derived vascular smooth muscle cells (VSMCs) and scRNA-seq approaches, we previously demonstrated that the 9p21.3 risk haplotype causally drives VSMC phenotypic plasticity, promoting an osteochondrogenic, calcification-prone state with impaired migratory capacity—features consistent with VSMCs observed in human atherosclerotic plaques^27^, and in line with previous reports observing increased calcification in plaque from 9p21.3 homozygous carriers^28^. While our study established a direct genotype-to-phenotype causal link between 9p21.3 and pathogenic VSMC behavior, it did not resolve the upstream regulatory mechanisms through which this non-coding locus exerts such broad effects on cell state.

Using a newly developed isogenic system that enables direct comparison of the risk and non-risk haplotypes within an identical genetic background, combined with long-read RNA sequencing, we uncover extensive, allele-specific remodeling of transcript expression and isoform usage across the VSMC transcriptome. We show that the 9p21.3 risk haplotype acts dominantly to reprogram RNA processing, shifting VSMCs toward a disease-prone osteochondrogenic state through predominantly trans-acting mechanisms.

At isoform-level resolution, we identify widespread haplotype-dependent splicing alterations affecting genes across multiple CAD-associated loci, revealing a previously unrecognized regulatory network, acting at different stages of atherosclerotic plaque formation, linking 9p21.3 to distal genomic regions. Among these, we identified the RNA helicase DDX5 (DEAD Box-Helicase 5) —linked to CAD risk through GWAS^29^ — as a key downstream effector whose haplotype-specific isoform usage modulates ANRIL expression, splicing factor abundance, and osteochondrogenic gene programs. Functional perturbation of DDX5 is sufficient to attenuate hallmark features of the risk-associated VSMC phenotype.

This work establishes alternative splicing as a central molecular conduit linking a major CAD risk locus to pathogenic vascular remodeling and redefines the mechanistic impact of the 9p21.3 risk locus in reshaping VSMC cell state primarily through isoform-level regulation of RNA processing rather than through conventional gene-expression changes. Together, these findings provide a high-resolution framework for interrogating how non-coding genetic variation drives cardiovascular disease.

## RESULTS

### RNA sequencing predicts splicing alterations in VSMCs driven by the 9p21.3 risk locus

In previous related work, we generated an isogenic human induced pluripotent stem cell (iPSC) panel for the 9p21.3 coronary artery disease (CAD) risk locus^26^. We then used bulk and scRNA sequencing to profile the transcriptome of terminally differentiated VSMCs across four genotypes: homozygous risk and non-risk haplotypes (RR and NN) and their isogenic knockouts (RRKO and NNKO)^26,27^. While this work uncovered large gene networks in vascular smooth muscle cells that drive acquisition of an osteochondrogenic, atherogenic-like cell state and identified transcriptional and functional dysregulation of adhesion, contraction, migration and calcification capacity, it did not describe upstream regulators responsible for these phenotypes. These key factors represent actionable genetics-based mechanisms and potential therapeutic targets. Hence, in the present study, we focused on identifying transcriptional regulators that exhibited modest but functionally consequential changes in expression, and thus more likely to reflect regulatory shifts (< absolute 1 log_2_ fold change). Indeed, Gene Ontology (GO) enrichment analysis using ClusterProfiler revealed significant upregulation of pathways associated with RNA splicing and mRNA processing in RR VSMCs (**Fig. 1A-B**, **Table S1A**), whereas pathways associated with ER and Golgi transport were reduced (**S1B, Table S1A-B**). These findings suggest that aberrant RNA splicing is a functional consequence of the 9p21.3 CAD risk haplotype.

**Figure 1.**
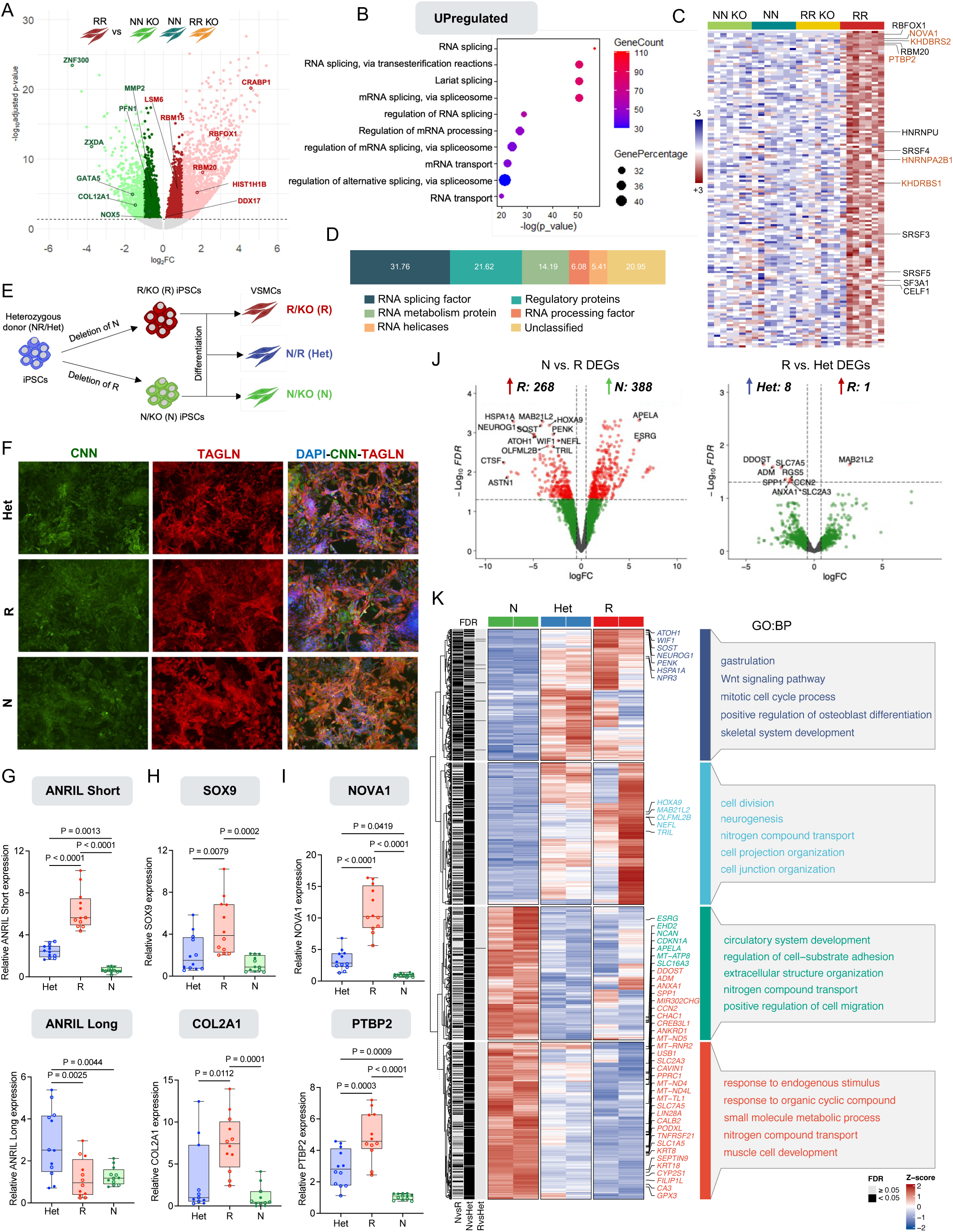
The 9p21.3 CAD risk locus impacts splicing processes. **(A)** Volcano plot of differentially expressed genes from bulk RNA sequencing performed on iPSC-derived VSMCs homozygous risk for the 9p21.3 locus (RR) compared to homozygous non-risk cells (NN) and isogenic VSMCs deleted of the 9p21.3 locus in both backgrounds (RRKO, NNKO) (Lo Sardo et.al, 2018). Light red and green are the genes analyzed in Lo Sardo et al. (2018). Red and green colored genes were analyzed in this study from absolute log_2_ fold change 0.1-1.0. **(B)** The top 10 gene ontology terms enriched in the differentially expressed genes in RR VSMCs between log_2_ fold change 0.1-1.0. **(C)** Heatmap of the top 149 differentially expressed splicing genes in iPSC-VSMCs across four genotypes at 9p21.3. N=7 per genotype. **(D)** Bar chart showing the biological function classification of the 149 splicing-related genes. **(E)** Schematic of experimental workflow for the generation of a new collection of isogenic iPSC. iPSCs are from one heterozygous donor (N/R- Het). Isogenic knockout lines were derived by TALEN-mediated haplotype editing to delete the entire CAD region and generate hemizygous lines (R/KO-R, N/KO-R). Cells were differentiated to VSMCs and used for long-read PacBio sequencing. **(F)** Immunocytochemistry for VSMCs markers for Het, R, and N VSMCs. Scale is 100 μm. **(G-I)** qPCR quantification of ANRIL short and long isoforms **(G)**, Osteochondrogenic markers SOX9 and COL2A1**(H)**, and the splicing genes NOVA1 and PTBP2 (**I)**. Data are from 2 independent cell lines per genotype and three independent VSMCs differentiation experiments. P-value calculated with One-way ANOVA with Bonferroni correction. **(J)** Volcano plots illustrate genome-wide differential gene expression between Non-risk (N) vs. Risk (R), and Risk vs. Heterozygous (Het) comparisons. logFC: log2 fold change. **(K)** Heatmap and Gene Ontology (GO) enrichment between N, Het, and R genotypes for DEGs.

Of the 477 genes associated with RNA metabolism and splicing processes, a striking 149 were upregulated in RR VSMCs relative to both non-risk and knockout lines (**Fig. 1C, S1C**). Several of these genes (e.g., RBFOX1^13^, RBM20^14^, SRSF3^15^) have established links to cardiovascular disease whereas others have not been described in connection with CAD (e.g., NOVA1, PTBP2, KHDBRS2). Importantly, these findings provide novel insight into the underlying pathology associated with the 9p21.3 risk haplotype.

Functional categorization of the 149-gene set revealed that close to a third (31.76%) encode canonical RNA splicing factors. The remainder include regulatory proteins (21.62% transcription factors, kinases, protein modifying enzymes), RNA metabolism factors (14.19%), RNA processing factors (6.08%), RNA helicases (5.41%) (**Fig. 1D**). Overall, this distribution support that a broad repertoire of upregulated RNA metabolism proteins - extending beyond the core spliceosome machinery - represents a distinct molecular signature of the 9p21.3 CAD risk haplotype.

### Establishing a system to compare 9p21.3 haplotype-specific contributions in VSMCs

To further characterize how 9p21.3 risk and non-risk haplotypes contribute to splicing-associated gene dysregulation, we generated a new isogenic iPSC lines collection to accurately dissect haplotype-specific effects within an otherwise identical genetic background. Starting with iPSC from a heterozygous individual, we performed targeted haplotype deletion to generate hemizygous clones retaining either the risk (R/KO or R) or non-risk (N/KO or N) allele (**Fig. 1E**). We validated genotypes by PCR and Sanger sequencing of SNPs rs1333049 and rs1057278 (**Fig. S1D-E**) and pluripotency using NANOG immunostaining (**Fig. S1F**). We then differentiated lines carrying only the risk allele (R), only the non-risk allele (N), or both alleles (Het) into VSMCs and verified successful differentiation for all lines by day 17 through immunostaining for the canonical VSMC markers calponin (CNN1) and transgelin (TAGLN), (**Fig. 1F**).

Consistent with prior studies^27^, qPCR analysis revealed a significant upregulation of short ANRIL isoforms (ending in exon 13) in R VSMCs compared with both N (p<0.0001) and Het controls (p<0.0001), while long ANRIL expression (ending in exon 20) did not differ between R and N (**Fig. 1G**). Similarly, expression of osteochondrogenic markers SOX9 and COL2A1 was significantly higher in R cells relative to N (SOX9 p=0.0002; COL2A1 p=0.0001) and Het cells (SOX9 p=0.0079; COL2A1 p=0.0112) (**Fig. 1H**). Additionally, we confirmed that R VSMCs maintain elevated expression of splicing-related genes, (e.g. NOVA1 and PTBP2) compared to N VSMCs (**Fig. 1I**), consistent with our initial analysis (**Fig. 1C**). Together, these results validate our newly generated isogenic lines and support that a single copy of the 9p21.3 risk haplotype is sufficient to reproduce the upregulation of short ANRIL isoforms, osteochondrogenic markers SOX9 and COL2A1, and the splicing-related genes observed in RR VSMCs.

### 9p21.3 allele-specific gene expression signature in VSMCs

We next employed PacBio long-read RNA sequencing to profile gene and transcript expression across our isogenic VSMCs lines. We achieved high-quality annotation for all libraries with comparable proportions across transcript classes (full splice matches 72.1%, incomplete splice matches 15.7%, novel in catalog 5.9% and novel not in catalog 5.6% **Fig S2A-C**). Leveraging the complex transcriptomic information allowed by the long-read sequencing we performed three different analysis: 1) overall gene expression to assess molecular signatures consistent with known 9p21.3-related cellular phenotypes; 2) transcript isoform expression to evaluate imbalance of specific isoforms of each gene, perhaps divergently mediated by the two alleles; 3) differential isoform usage, to assess allele-specific isoform composition imbalance even when overall gene expression is not compromised overall.

We performed differential gene expression (DEG) analyses across three pairwise comparisons (N vs R; N vs Het; R vs Het). Comparing N and R VSMCs identified 656 DEGs (p < 0.05), with 268 genes upregulated and 388 downregulated in R VSMCs (**Fig 1J**). The N vs Het comparison identified 1,324 DEGs (p < 0.05), with 645 genes upregulated and 679 downregulated in Het VSMCs (**Fig S2D**). In contrast, R and Het VSMCs were highly similar, differing by only nine DEGs, indicating near complete transcriptional convergence (**Fig. 1J**). Notably, N vs R and N vs Het shared 495 DEGs (**Fig S2E**), demonstrating that the presence of a single risk allele is sufficient to impose the risk-associated transcriptional program.

Gene Ontology (GO) enrichment analysis of DEGs from N vs R and N vs Het comparisons revealed highly similar biological signatures, consistent with our previous findings in an independent isogenic line collection^26,27^. Genes downregulated in R VSMCs were enriched for pathways related to vascular development, wound healing, and smooth muscle maintenance, including circulatory system development, cell adhesion, migration and muscle development (**Fig 1K**). Among the most strongly downregulated genes in both R and Het VSMCs was APELA/ELABELA (**Fig 1J, S2D**), an endogenous ligand of the Apelin receptor with established roles in heart development and coronary vessel sprouting ^30,31^. This suggests an undescribed role for the 9p21.3 locus in coronary vessel maintenance, vascular remodeling, and angiogenesis — processes that may be compromised by the presence of the risk allele.

In contrast, genes upregulated in R VSMCs were predominantly associated with skeletal development, osteoblast differentiation, Wnt signaling pathway and cell-cycle regulation (**Fig 1K**). The N vs Het comparison closely mirrored these patterns with genes upregulated in Het cells similarly enriched for osteoblast differentiation pathways (**Fig 1K**).

Together, these results reinforce that the 9p21.3 risk haplotype drives a robust shift towards osteochondrogenic programs in VSMCs. This effect is independent of genetic background: the presence of the single risk allele at 9p21.3, whether in hemizygous or heterozygous form, is sufficient to redirect VSMCs away from vascular maintenance programs toward an osteogenic differentiation trajectory. Within a shared genetic background, selective retention of the risk or non-risk haplotype thus biases the global transcriptomic landscape toward distinct cell fates— vascular versus osteogenic —highlighting a central role for 9p21.3 in VSMC fate determination.

### Haplotype-dependent regulation of transcript isoforms shapes VSMC cell state

Because long-read sequencing resolves full-length transcripts, it enables the detection of isoform-level differences that are often missed by short-read approaches. Leveraging the PacBio long-read RNA sequencing, we profiled isogenic VSMCs carrying distinct 9p21.3 haplotypes and systematically assessed haplotype-dependent transcript regulation.

Across the same three pairwise comparisons (N vs R, N vs Het, and R vs Het), we identified extensive allele-dependent transcript differences. In the N vs R comparison, 754 transcripts were upregulated and 751 downregulated in R VSMCs. Similarly, the N vs Het comparison identified 775 transcripts upregulated and 579 downregulated in Het relative to N VSMCs (**Fig 2A**). These two comparisons shared 734 differentially expressed transcripts (**Fig S2F-G**), indicating that the presence of the risk allele drives a largely shared transcriptomic program in both hemizygous and heterozygous states.

**Figure 2.**
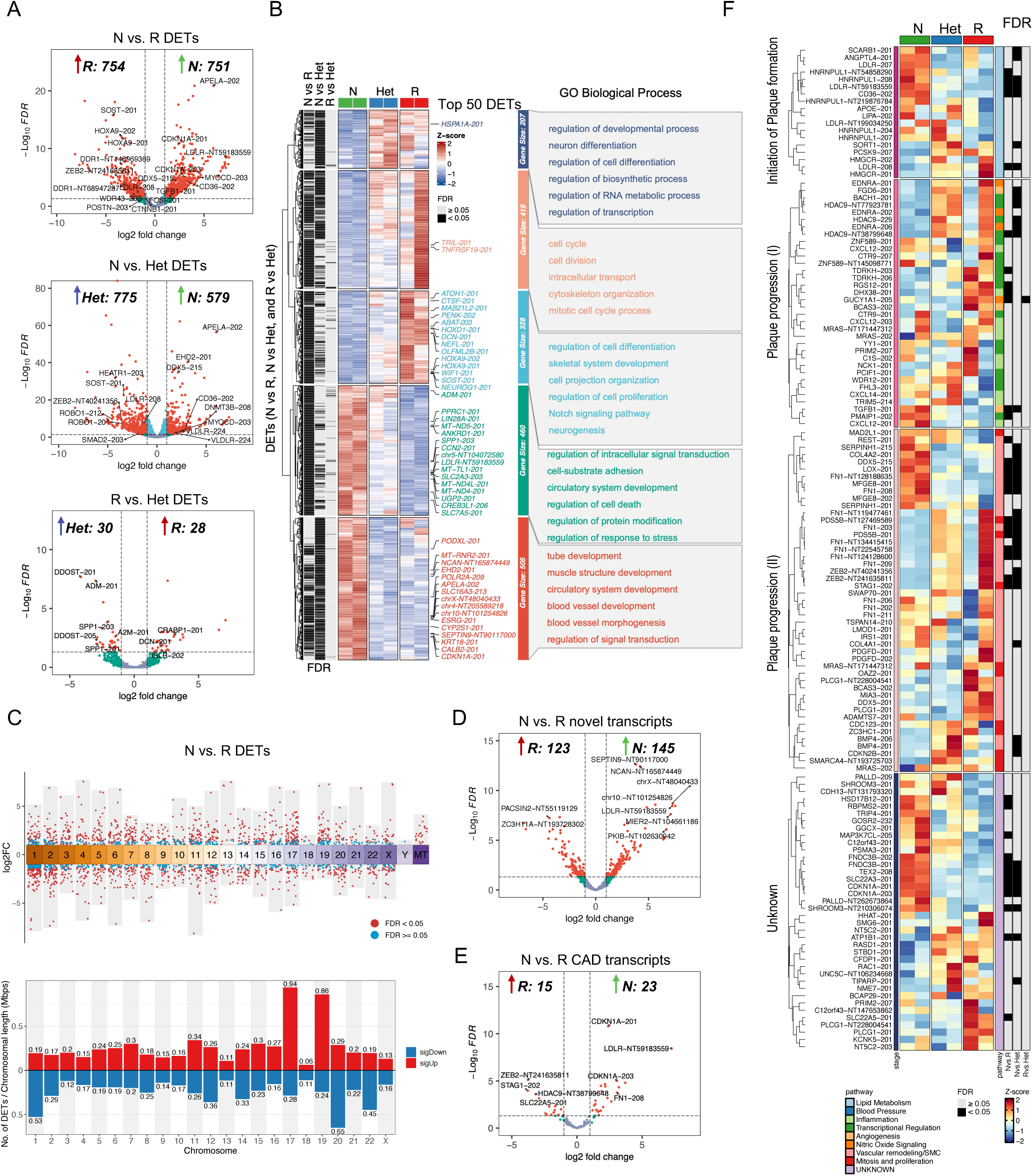
Haplotype-biased transcript isoforms expression controls cell fate in VSMCs. **(A)** Volcano plots illustrate genome-wide differential transcript expression between Non-risk (N) vs. Risk (R), Non-risk vs. Heterozygous (Het), and Risk vs. Heterozygous comparisons. **(B)** Heatmap and Gene Ontology (GO) enrichment between N, R, and Het genotypes for DETs upregulated and downregulated. **(C)** Chromosomal distribution of DETs with log_2_ fold changes across the genome for N vs R haplotypes. Each dot represents a transcript positioned along chromosomes 1–22, X, and Y, as well as mitochondrial DNA. Bar plots showing the number of DETs per chromosome normalized to chromosome length for N vs R. Red bars indicate upregulated DETs, blue bars indicate downregulated DETs. **(D)** Volcano plots showing differential expression of novel transcripts (NIC and NNIC), for N vs R VSMCs. **(E)** Volcano plots showing differential expression of CAD-associated transcripts, for N vs R VSMCs. **(F)** Heatmap enrichment between N, R, and Het genotypes for DETs related to genes associated to CAD progression according to Kesseler et.al, 2019.

Notably, although R and Het VSMCs differed by only nine genes at the gene-expression level, long-read sequencing detected 58 differentially expressed transcripts between these genotypes (**Fig 2A**). These data highlight the increased sensitivity of full-length transcriptome profiling in detecting subtle but biologically relevant isoform-level differences.

Transcripts differentially expressed in N vs R comparison mapped to 1268 unique genes of which ∼71% were also altered in the N vs Het comparison, indicating strong transcriptional convergence between R and Het VSMCs. This overlap further supports a dominant effect of the risk haplotype on transcript-level regulation.

Unsupervised clustering of differentially expressed transcripts identified five major expression patterns (**Fig. 2B**). Clusters I and II comprised transcripts upregulated in both R and Het cells and were enriched for pathways related to cell differentiation, RNA metabolic processes, cell-cycle regulation and cytoskeletal organization. Cluster III contained transcripts uniquely upregulated in R cells, associated with skeletal development, neuronal projection, and Notch signaling. In contrast, Clusters IV and V represented non-risk-associated programs: transcripts upregulated in N VSMCs (and partially retained in Het VSMCs), enriched for proliferation and circulatory system development (Cluster IV), and a distinct N-specific cluster absent in risk-bearing cells enriched for tube morphogenesis, blood vessel formation and vascular structure organization (Cluster V) (**Fig 2B**).

To assess whether haplotype-dependent transcript changes localized to specific genomic regions, we examined their chromosomal distribution across comparisons. In both N vs R and N vs Het, allele-specific dependent transcript changes were broadly distributed across the genome (**Fig 2C and S2H**). After normalization for chromosomal length, we observed enrichment of upregulated transcripts on chromosomes 17 and 19 and of downregulated transcripts on chromosome 20 (**Fig. 2C and S2H**). In contrast, transcripts differentially expressed between R and Het did not show chromosomal enrichment. Importantly, we detected no enrichment on chromosome 9, consistent with a predominantly trans-acting mechanism of 9p21.3-mediated regulation rather than local cis effects (**Fig 2C)**.

Long-read sequencing further enabled the identification of hundreds of previously un-annotated transcripts isoforms. In total, we detected 268 novel transcripts (NIC-Novel In Catalog and NNIC- Novel Non In Catalog; p<0.05), corresponding to 248 genes, differentially expressed in R versus N VSMCs (**Fig 2D**), and 382 novel transcripts (331 genes) in Het versus N cells, with 141 shared between these two comparisons (**Fig S2I**). Gene ontology analysis linked these novel isoforms primarily to pathways regulating cell differentiation. Together, these findings reveal extensive, previously uncharacterized isoform diversity associated with the 9p21.3 risk haplotype that may contribute to VSMC cell-state transitions.

### 9p21.3 drives transcript and isoform remodeling across CAD loci through predominantly trans-acting mechanisms

To assess the effect of the two haplotypes at 9p21.3 on other CAD-related genes, we examined 198 genes predicted to be causal across all known CAD-loci^8^. Transcript-level analysis identified 32 differentially expressed transcripts in the N vs R comparison and 37 transcripts in the N vs Het comparison (**Fig 2E, S2J**). This analysis confirmed previously reported expression changes detected by bulk and scRNAseq (e.g. CXCL12, COL4A1, COL4A2, REST and others)^26,27^ and additionally identified specific annotated isoforms and previously unannotated transcripts that were not resolved in earlier studies. Notably, our isogenic long-read approach uncovered changes in several CAD genes not previously detected, including genes involved in lipid metabolism-associated genes (ANGPTL4, PCSK9, SCARB1); vascular remodeling (TGFB1, FNDC3B); macrophage activation (TEX2); proliferation and dedifferentiation (CDKN1A, MFGE8) and gene expression regulation (STAG1) in R or Het compared to N VSMCs (**Fig 2F, S2K**).

Importantly, isoform-level analysis revealed additional haplotype-dependent alterations in transcripts of LDLR (related to lipid metabolism), LMOD1 and EDNRA (involved in VSMC dedifferentiation), HNRNPUL1 and DDX5 (associated with RNA processing) (**Fig 2F)**. Classification of the 198 CAD genes based on predicted biological function and stage of atherosclerotic plaque development revealed enrichment among genes involved in plaque initiation and progression, although significant transcript alterations were observed across all stages of plaque formation (**Fig. 2F**). These findings support a model in which the 9p21.3 risk haplotype contributes to multiple phases of CAD pathogenesis through widespread transcript and isoform remodeling at distal genomic loci.

Work from our group and others has shown that the 9p21.3 locus alters the expression of its neighboring genes, CDKN2A and CDKN2B, although substantial variability has been reported, consistent with tissue-and cell type-specific effects^26,27,32^. We sought to investigate local splicing alterations and isoform-switching events affecting genes on chromosome 9. Our initial analyses did not identify a significant enrichment of splicing events on this chromosome, suggesting that the local impact of the 9p21.3 haplotype on splicing is modest (**Fig S3A**). Together, these results indicate that the 9p21.3 risk haplotype reshapes RNA processing predominantly through trans-acting effects at distal CAD loci, rather than through extensive local splicing regulation.

### Haplotype-specific isoform usage reveals novel targets of 9p21.3 in VSMCs

The isoform diversity revealed by long-read sequencing enabled the detection of transcript-level regulatory changes occurring independently of alterations in total gene expression. This approach allowed the identification of differential transcript usage (DTU) even in cases where overall gene expression remained comparable across genotypes. Because individual genes can generate multiple transcript isoforms with distinct functional properties, changes in relative isoforms abundance can substantially influence cellular behavior and downstream regulatory programs^33^. DTU analysis revealed extensive haplotype-dependent isoform regulation associated with the 9p21.3 locus. Specifically, 187 transcripts corresponding to 132 genes exhibited differential isoform usage in the N vs R comparison, while 91 transcripts corresponding to 66 genes were altered in the N vs Het comparison (**Fig. S4A; Fig. 3A**). In contrast, the R vs Het comparison yielded only two genes with significant DTU, consistent with the minimal transcriptional differences observed between these genotypes at the gene-expression level (**Fig 3A**).

**Figure 3.**
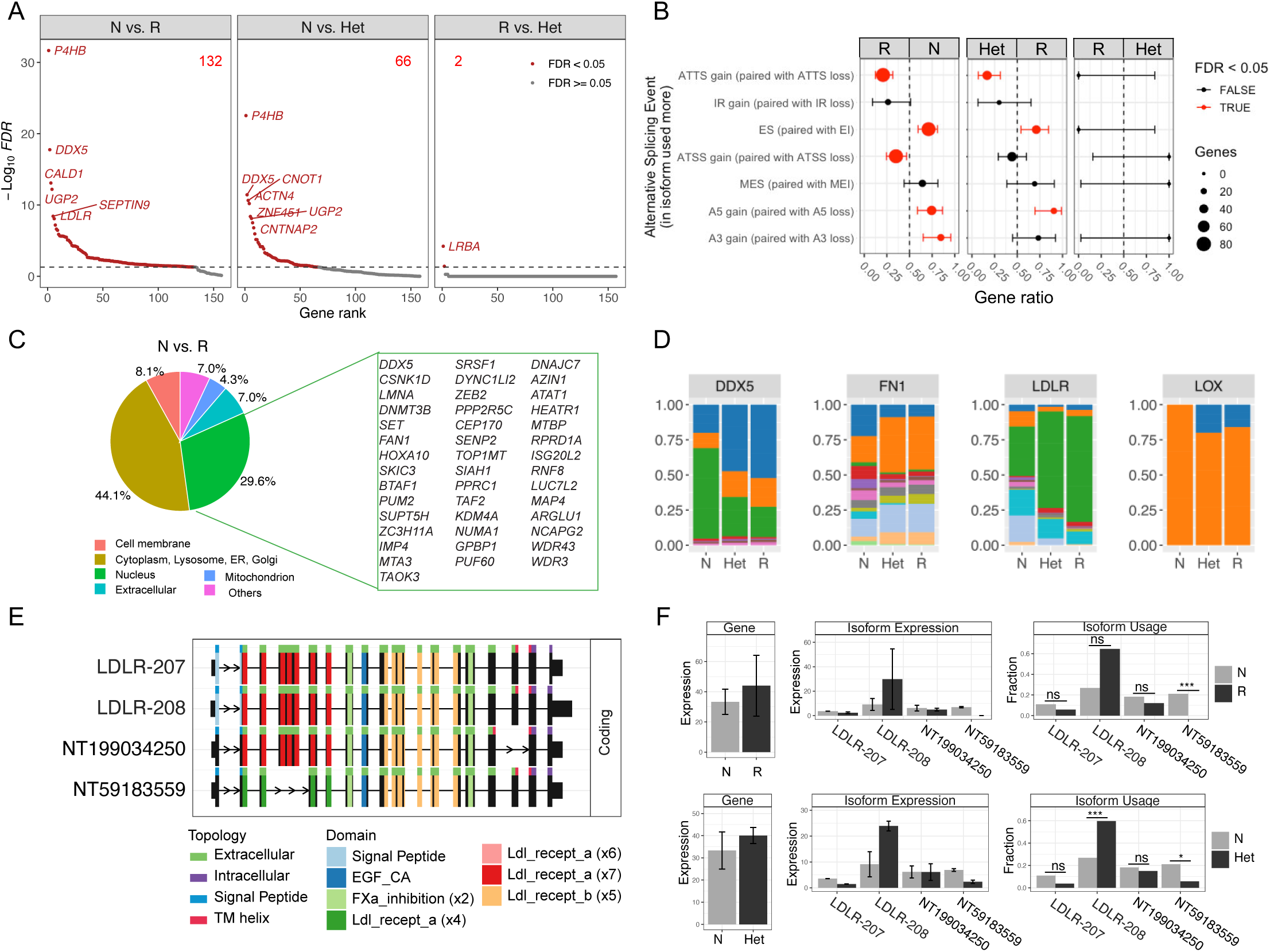
The risk haplotype at 9p21.3 drives divergent transcript isoform usage in VSMCs. **(A)** Ranked gene plots showing differential isoform usage between Non-risk (N), Risk (R), and Heterozygous (Het) groups. Red dots indicate genes with significant changes (FDR < 0.05). **(B)** Alternative splicing event types associated with isoforms used more across comparisons. Each dot represents a splicing event category, with dot size indicating the number of affected genes. Significant events (FDR < 0.05) are shown in red. **(C)** Pie charts indicate the cellular localization of proteins encoded by genes undergoing significant isoform switching between N vs. R haplotypes. **(D)** Stacked bar charts illustrating isoform usage proportions of key CAD-related genes (DDX5, FN1, LDLR, LOX) across N, Het, and R genotypes. Each color represents a distinct transcript isoform. **(E)** Schematic representation showing distinct exon composition and domain architecture of LDLR protein isoforms. **(F)** Bar graphs show gene-level expression, isoform-specific expression, and isoform usage of LDLR transcripts between N vs. R, and N vs. Het.

The N vs R analysis uncovered isoform usage alteration in several cardiovascular-related genes not detected in previous studies^26,27^, including P4HB, DDX5, CALD1, UGP2, SEPTIN9 and LDLR (**Fig S4E-F**). Additionally, 41 genes displayed shared isoform usage changes in both R and Het VSMCs compared with N VSMCs (**Fig S4B**), further supporting a dominant effect of the risk haplotype on isoform regulation.

To characterize the splicing events underlying these isoform changes, we classified alternative splicing events across comparisons, including Alternative Transcription Termination Sites (ATTS), Intron Retention (IR), Exon Skipping (ES), Alternative Transcription Start Sites (ATSS), Multiple Exon Skipping (MES), Alternative 5′ Splice Sites (A5 gain), and Alternative 3′ Splice Sites (A3 gain). Isoforms preferentially expressed in R VSMCs showed significant enrichment for ATTS and ATSS events relative to N cells, whereas N VSMCs preferentially utilized isoforms arising from ES, A5, and A3 events (**Fig 3B, S4C**). Het cells closely mirrored the R pattern, with reduced ES and A5 usage and increased ATTS relative to N. No significant differences were detected between R and Het cells.

We next assessed the predicted subcellular localization of proteins encoded by differentially used transcripts. In the N vs R comparison, 44.1% localized to the cytoplasm or cytoplasmic organelles (including lysosome, ER, and Golgi), 29.6% to the nucleus, 8.1% to the cell membrane, 7% extracellularly, and 4.3% to mitochondria (**Fig 3C**). Similar distributions were observed in the N vs Het comparison (**Fig S4D**). Given that many cytoplasmic and membrane-localized proteins are involved in downstream phenotypic processes such as cell morphology and ECM production, we focused subsequent analysis on transcripts encoding nuclear-localized proteins to identify potential upstream regulatory factors.

Among nuclear-localized DTU targets in the N vs R comparison, we identified transcription factors and cofactors (DDX5, TAF2, PPRC1, MTA3, HOXA10, ZEB2, BTAF1, GPBP1), RNA metabolism and splicing regulators (HEATR1, PUM2, DDX5, ZC3H11A, PUF60, SRSF1, IMP4), RNA binding proteins (LUC7L2, WDR43, WR3) and chromatin associated enzymes (DNMT3B, SET, KDM4A, LMNA) (**Fig 3C**). These findings suggest that haplotype-specific isoform regulation at 9p21.3 affects multiple layers of gene regulation in VSMCs, including transcriptional and RNA-processing machinery. Chromosome 9–focused DTU analysis identified limited local effects (IL11RA, SET, UNC13B, and SLC25A25 in N vs. R, NCBP1, MED22 in N vs. Het), with no significant DTU detected between R and Het cells (**Fig S3B–D**).

DTU analysis of CAD-GWAS genes further highlighted DDX5, LDLR, LOX, and FN1 as prominent haplotype-sensitive targets (**Fig 3D, Fig S4E-H**). Notably, four distinct coding isoforms of LDLR were detected with imbalanced usage across genotypes (**Fig 3E-F**). Isoform LDLR-208 was significantly enriched in Het VSMCs and showed a similar trend in R VSMCs, whereas a novel isoform identified in our dataset (NT59183559) was markedly depleted in R and Het cells (**Fig 3F**). The remaining two isoforms (LDLR-207 and NT199034250) were comparably expressed across genotypes (**Fig 3E-F**).

Collectively, these findings demonstrate that the 9p21.3 risk haplotype drives extensive isoform-level remodeling of transcriptional regulators and CAD-associated genes, frequently without corresponding changes in total gene expression.

### The helicase DDX5 is a novel downstream effector of the 9p21.3 risk locus

Two independent analyses—DTU profiling and CAD-gene transcript analysis—identified the DEAD-box RNA helicase DDX5 as a prominent target of haplotype–specific regulation at the 9p21.3 locus. DDX5 has previously been implicated in cardiac function and heart failure^34^, prompting us to investigate its role in the 9p21.3-dependent VSMC phenotype. Although overall DDX5 gene expression did not differ significantly across genotypes, isoform-level analysis revealed marked haplotype-specific differences. Among the three major DDX5 isoforms — two coding isoforms (DDX5-201 and DDX5-201-0) and one intron-retained isoform (DDX5-215), predicted to be NMD-sensitive — R and Het VSMCs preferentially expressed the coding isoform DDX5-201 and showed a significant reduction of the DDX5-215 isoform (**Fig. 4A-B**). Isoform-specific qPCR confirmed increased DDX5-201 expression in R and Het VSMCs relative N cells (**Fig 4C**).

**Figure 4.**
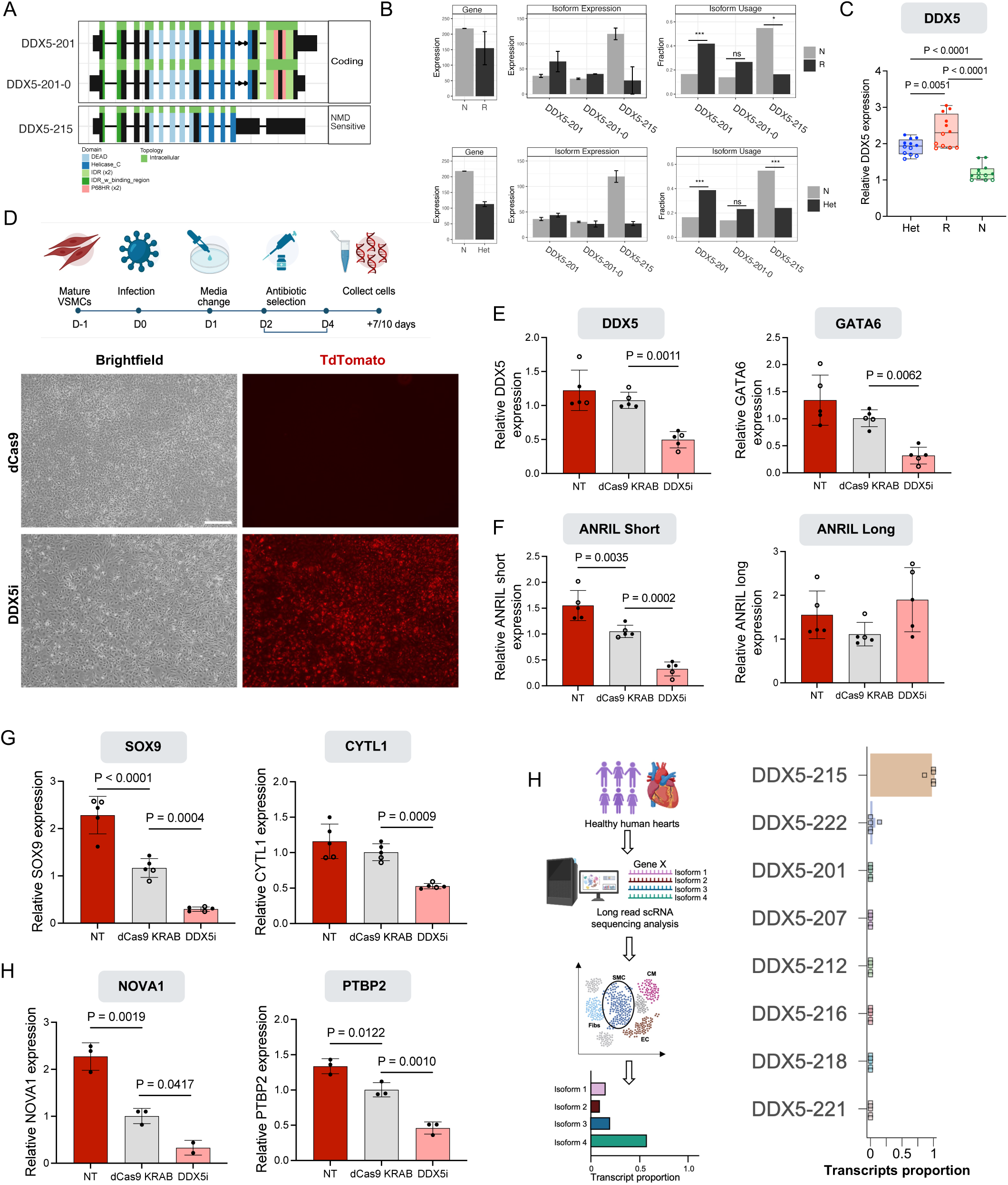
DDX5 targeting ameliorate the effect of the 9p21.3 risk haplotype in VSMCs. **(A)** Schematic representation showing distinct exon composition and domain architecture of DDX5 protein isoforms. **(B)** Bar graphs show gene-level expression, isoform-specific expression, and isoform usage of DDX5 transcripts between N vs. R, and N vs. Het. **(C)** qPCR quantification of DDX5 isoform using isoform-specific primers. Data are from 2 independent cell lines per genotype and three independent differentiation experiments. **(D)** Schematic of CRISPR interference (CRISPRi) timeline in mature Risk VSMCs. Representative brightfield and fluorescence images of VSMCs upon CRISPRi with dCas9-KRAB only (control) and DDX5i- TdTomato (transduced with dCas9-KRAB and DDX5-specific gRNA with TdTomato reporter). Scale is 200 μm. **(E-F)** qPCR quantification of DDX5, and GATA6 **(E)** and ANRIL isoforms **(F)**. **(G-H)** qPCR quantification of osteochondrogenic markers **(G)**, and splicing genes **(H)**. Data are from two independent experiments. P-value calculated with One-way ANOVA with Bonferroni correction. **(I)** Detection of DDX5 transcripts(reference-based) in the Heart Isoform Atlas (https://gaolabtools.shinyapps.io/heart-isoform-atlas/) from Pan et al 2025. The quantification relates to the smooth muscle cell population in non-disease heart samples.

To test whether altered DDX5 isoform composition contributes functionally to the risk-associated phenotype, we performed CRISPR interference (CRISPRi) in iPSC-derived R VSMCs, which predominantly express DDX5-201. Cells were transduced with a dual-vector system expressing dCas9-KRAB and a DDX5-targeting guide RNA with TdTomato expression used to monitor transduction efficiency (**Fig. 4D-E, S5A**). Effective DDX5-201 knockdown (as measured by our isoforms-biased qPCR) resulted in reduced expression of GATA6, a known downstream target, confirming functional suppression (**Fig. 4E**). Notably, DDX5 knockdown let to a marked reduction in short ANRIL isoforms, while long ANRIL isoforms remained unchanged (**Fig. 4F**). Expression of osteochondrogenic markers SOX9 and CYTL1 was also significantly reduced (**Fig. 4G**). In addition, expression of splicing factors identified in the discovery analysis (NOVA1 and PTBP2) (**Fig 1C**), decreased following DDX5 suppression (**Fig. 4H**). In contrast, overexpression of NOVA1 or PTBP2 in N VSMCs did not alter DDX5 expression (**Fig. S5B–C**), supporting a model in which DDX5 functions upstream of broader splicing alterations.

To assess translational relevance, we examined DDX5 isoform expression in a single-cell atlas of splicing isoforms from healthy human hearts^35^. In this dataset, the intron-retained DDX5-215 isoform was the most predominantly expressed in smooth muscle cells. Consistently, DDX5-215 was the most abundant isoform in N VSMCs in our system (**Fig. 4I**). In contrast, R VSMCs exhibited a pronounced reduction of DDX5-215 accompanied by increased expression of DDX5-201, an isoform expressed at low levels in healthy heart tissue, suggesting an aberrant CAD-associated isoform signature.

Collectively, these findings identify DDX5 isoform regulation as a downstream effect of the 9p21.3 risk haplotype and link allele-specific RNA-processing changes to VSMC phenotypic modulation.

## Discussion

Alternative splicing is increasingly recognized as a central pathogenic mechanism in cardiovascular disease, contributing to atherosclerosis^9–11^, heart failure^13,14^, and vascular remodeling^12^. Variants disrupting splicing regulators or splice sites have been implicated in congenital heart disease, impaired cardiac or vascular function^13–16^. However, a key unresolved question is whether common non-coding CAD risk loci exert dominant effects on vascular cell state and disease susceptibility by perturbing RNA processing—not only through altered gene expression, but also through more subtle mechanisms such as transcript isoform imbalance in the absence of major changes in total gene expression. Here, we address this question by investigating RNA processing dysregulation at the strongest known CAD locus, the 9p21.3. The 9p21.3 coronary artery disease (CAD) risk locus has been extensively replicated across multiple GWAS as a major genomic region associated with increased CAD risk in diverse human populations^17–19,21,36,37^. Although identified nearly two decades ago, the functional mechanisms of this locus remain challenging to decipher. Over the past decade, our group and others have sought to unravel this genomic enigma using a range of experimental approaches^24,38–43^. Modeling this locus in mice has proven particularly difficult, as the syntenic region lacks the characteristic human haploblock of SNPs and an accurately annotated homolog of ANRIL, the lncRNA encoded within the human locus^24,25^.

To overcome these limitations, our group previously generated a human-specific knockout model of the 9p21.3 locus using induced pluripotent stem cells (iPSC) and haplotype-specific genome editing. Using this system, we established a causal genotype-to-phenotype relationship between the 9p21.3 CAD risk locus and vascular smooth muscle cell (VSMC) behavior^26^. Specifically, we demonstrated that the risk haplotype alters VSMC plasticity, promoting acquisition of an osteochondrogenic transcriptional program accompanied by increased calcification propensity and reduced migratory capacity^27^.

While these studies established a causal role for 9p21.3 in vascular calcification—a defining feature of advanced atherosclerotic plaques—the distinct contributions of the risk and non-risk haplotypes in driving divergent VSMC states remained incompletely understood. To address this gap, we developed a platform combining haplotype-biased genome editing with long-read RNA sequencing, enabling transcriptome-wide analysis of RNA regulation at isoform-level resolution. This approach allowed us to directly compare the effects of the two major 9p21.3 haplotypes within an isogenic background and revealed that this locus shapes VSMC phenotypic state through widespread reprogramming of alternative splicing across the transcriptome.

Our findings demonstrate that the 9p21.3 risk haplotype exerts a dominant effect on VSMC phenotypes over the non-risk allele *in vitro*, consistent with the elevated disease risk observed in heterozygous individuals^17–19^. Notably, the risk-associated molecular signature persists in heterozygous settings, indicating that a single copy of the risk allele is sufficient to impose a disease-prone transcriptional and splicing program. To our knowledge, this is the first isogenic human system that enables direct comparison of how the two distinct 9p21.3 haplotypes determine VSMC cell-state programs under both hemizygous and heterozygous conditions. Although modest, we also observed transcriptional differences between VSMCs carrying only the risk haplotype and those carrying both risk and non-risk haplotypes, suggesting that haplotype dosage may fine-tune downstream regulatory programs.

Importantly, our study provides the first direct evidence linking alternative splicing regulation to the 9p21.3 CAD risk locus. We show that a common CAD risk haplotype can dominantly reprogram transcript isoform expression and usage across the VSMC transcriptome, often without corresponding changes in overall gene expression. These findings highlight RNA isoform regulation as a critical yet underappreciated mechanism through which non-coding genetic variation can influence vascular cell behavior and disease susceptibility.

Our new approach identified divergent effects of the 9p21.3 risk and non-risk haplotypes on expression of APELA (ELA/ELABELA) in VSMCs. APELA is an endogenous ligand of the Apelin receptor (APLNR) and plays key roles in coronary artery morphogenesis, vascular branching, and angiogenic remodeling after injury^31,44,45^. The allele-specific modulation of APELA by the 9p21.3 haplotype suggests a mechanistic link between this CAD risk locus and coronary artery remodeling capacity, potentially influencing vascular repair and adaptive responses under stress.

By integrating our isoform-level data with the most recent catalog of CAD-associated loci and putative causal genes, we uncovered a network linking the 9p21.3 locus to isoform remodeling across multiple CAD-relevant genes, including LDLR, ANGPTL4, and PCSK9, major pharmacological targets of CAD treatment. These connections, not previously described, suggest that 9p21.3 operates within a broader regulatory landscape that intersects with other major genetic risk factors for CAD and acts at different stages of atherosclerotic plaque formation, perhaps explaining the strong impact of this genomic locus in CAD risk. The widespread isoform remodeling observed across distal CAD loci, coupled with the absence of local enrichment on chromosome 9, supports a predominantly trans-acting model for 9p21.3-mediated regulation.

The present study provides the first long-read RNA sequencing dataset of cells carrying distinct 9p21.3 haplotypes and identified thousands of differentially expressed transcripts—far exceeding those detected by previous short-read approaches— including both NIC (Novel In Catalog) or NNIC (Novel Not in Catalog) transcripts, associated with allele-specific regulation. These previously unannotated isoforms revealed a previously inaccessible molecular signature of this locus and, more broadly, expand the transcriptomic complexity of VSMCs to be leveraged as a valuable resource for future functional studies aimed at understanding cell-state transitions and regulatory control in vascular disease.

A key strength of our controlled isogenic system is the ability to identify genes whose overall expression remains stable across haplotypes but that undergo profound isoform switching. This reveals a mode of regulation that is invisible to conventional gene-level analysis and underscores the importance of transcript usage and RNA processing in mediating CAD risk.

Among downstream targets identified, DEAD-box helicase 5 (DDX5) emerged as a prominent effector of haplotype-specific regulation of the 9p21.3 locus. Using CRISPR interference, we demonstrate that modulation of the risk-associated DDX5 isoform in risk VSMCs reduces ANRIL short isoform expression, the osteochondrogenic transcriptional signature and other splicing regulators associated with the risk haplotype. Although DDX5 has been implicated in heart failure^34^ and vascular remodeling^46^, its role in CAD predisposition and CAD-related VSMC phenotypic modulation has not previously been explored. Our findings establish a causal link between 9p21.3 and DDX5 and position this RNA helicase as a key mediator of splicing alterations in VSMC plasticity.

Consistent with these findings, we show that VSMCs derived from non-risk iPSCs exhibit an isoform signature closely resembling that of healthy human heart smooth muscle cells (DDX5-215-biased), as defined by a recent single-cell isoform atlas^35^. In contrast, risk haplotype-bearing VSMCs display depletion of the DDX5-215 isoform and enrichment of the DDX5-201 isoform, which is minimally expressed in healthy heart tissue. This aberrant isoform balance suggests a CAD-associated RNA-processing signature driven by the 9p21.3 risk haplotype. We speculate that the 9p21.3 haplotypes may influence DDX5 splicing and expression through long-range chromatin interactions, a hypothesis supported by the enrichment of differential transcript expression on chromosomes 17 (where DDX5 is located).

In summary, we present a high-resolution dissection of how the 9p21.3 CAD locus in VSMCs shapes vascular smooth muscle cell state through dominant, haplotype-specific, and predominantly trans-acting regulation of RNA isoform usage. By integrating isogenic haplotype editing with long-read transcriptomics, this work reveals a previously inaccessible layer of gene regulation linking non-coding genetic risk to VSMC phenotypic plasticity and identifies DDX5 as a key downstream effector of these programs. These findings suggest that isoform-level regulation represents a generalizable mechanism through which common CAD risk loci influence vascular disease susceptibility.

## Materials & Methods

### Data availability

Long-read bulk RNA sequencing datasets will be publicly available as of the date of publication. All other data supporting the findings in this study are included in the main article and associated Supplementary materials. Unedited iPSC lines SCRP2411i (Cell Line Alias: KBET2411i) (RRID: CVCL_JV64) are available at WiCell in the NHLBI Next Gen - coronary artery disease and myocardial infarction (Dr. Eric Topol, Scripps Research Institute) collection.

### Ethical compliance and cell lines

The authors have complied with all ethical regulations. All iPSC lines used in this study were previously derived and characterized under a study approved by the Scripps Research Institute IRB (IRB#115676, IRB#07-4714)^47^. The cells were transferred and maintained under a study approved by the University of Wisconsin-Madison through a Material Transfer Agreement (MSN246500). The iPSC lines included in this study were generated from Peripheral Blood Mononuclear cells (PBMCs) from blood donors consented under the IRB#115676, IRB#07-4714. All iPSC lines used derive from male donors: HE554 (SCRP2411i; Cell Line Alias: KBET2411i) (RRID: CVCL_JV64); C512 (SCRP0517i; Cell Line Alias: KBET0517i) (RRID: CVCL_JV34); and HE463 (SCRP2307i; Cell Line Alias: KBET2307i) (RRID: CVCL_JV60). The choice to restrict this study to male donors was made to ensure line stability in relation to X inactivation, which can spontaneously change in female iPSC lines. Our study was not designed to detect sex-based phenotypes, and sex was not a determinant in our experiments. We have based our decision to exclude sex from consideration on the CAD GWAS data available thus far, which shows no compelling evidence of sex differences in 9p21.3-based risk of CAD.

### Published dataset analysis

The dataset from Lo Sardo et al. is available GEO: GSE120099

<u>Heatmaps</u>: Heatmaps of published data were generated using the heatmap.2 command from the ggplot2 package (v3.5.1).

<u>Volcano plots</u>: Volcano plots of published data were generated using the ggplot command, also from the ggplot2 package.

<u>Gene Ontology (GO) Enrichment Analysis</u>: GO analysis was conducted using the clusterProfiler package (v3.0.4, RRID:SCR_016884).

### iPSC derivation, differentiation and maintenance

iPSCs were cultured on Matrigel-coated (Corning 354277) cell culture plates and mTeSR (Stem Cell Technologies cat. 85850), cultured at 37°C under normoxic conditions (20% O_2_) with 5% CO_2_. iPSCs were differentiated into VSMCs as previously described (Sardo et al. 2018; Cheung et al. 2014). VSMCs were maintained on VSMC media (DMEM, Gibco 11965092; 10% BenchMark™ Fetal Bovine Serum, Gemini; Glutamax, Gibco 35050061; Penicillin/Streptomycin, Gibco 15140122; MEM Non-Essential Amino Acids, Gibco 11140050).

### Generation of hemizygous iPSC

Isogenic hemizygous lines were generated from a male heterozygous donor at the 9p21.3 locus (carrying one risk and one non-risk allele) by TALEN-mediated haplotype editing as we previously described and validated. After single cell subcloning and iPSC clone expansion, genotyping was performed by primer sequences including: GJC 344F (5′ CATACAGGTCCCTGGCACTAA 3′), GJC 346F (5′ CGAAGGGCTTCCCTGTCTA 3′), and 347R (5′ GACTTTCCCCCA CAATGAAA 3′). Genotyping PCR was performed using Q5 Hot Start DNA <u>polymerase</u> (NEB M0493S) in 25uL reactions. Reaction parameters include: 30 cycles, 58 °C annealing, 30sec extension, 200ng genomic DNA template per reaction, and primer concentration of 200nM each. Based on genotyping, we selected iPSC clones with deletion of either the risk or non-risk 9p21.3 locus, confirmed by Sanger sequencing for SNPs rs1333049 and rs10757278.

### Long-read bulk RNA sequencing

Total RNA samples submitted to the University of Wisconsin-Madison Biotechnology Center Gene Expression Center (*RRID:SCR_017757)* were assayed on NanoDropOne Spectrophotometer and Agilent 4200Tapestation to assess purity and integrity. RNA concentrations were accurately quantified by Qubit 4.0. RNA samples with RIN > 9 were selected for library construction. 300ng total RNA was converted to single-stranded cDNA and amplified using the Iso-Seq Express 2.0 Kit (Pacific Biosciences). Iso-Seq libraries were constructed with the SMRTbell Template Prep Kit 3.0 (Pacific Biosciences). Library quantity and quality were assessed by Qubit 4.0 and Agilent 4200Tapestation, respectively. Libraries were sequenced on one PacBio Revio SMRT cell. Revio polymerase kit v1.0 and Revio SMRT cell v1.0 used (Pacific Biosciences). Library prep and sequencing were conducted at the Gene Expression Core of the Biotechnology Center at UW-Madison.

### Bioinformatic analysis of long-read bulk RNA sequencing

All codes and scripts used in this study will be available on GitHub.

### Isoform Discovery and Alignment

We identified and characterized gene isoforms from PacBio HiFi reads using the IsoSeq v4.3 pipeline. First, the initial processing of the raw BAM files began with the *lima* tool to identify and trim the 5’ and 3’ cDNA primers. This step also used the embedded barcode sequences to demultiplex the pooled sequencing data into sample-specific files. These full-length reads then underwent further cleaning with *isoseq refine*. This step identifies and trims the 3’ poly(A) tail and detects and splits artificial concatemers—ligation artifacts where multiple distinct cDNA molecules are sequenced as a single long read. The resulting full-length, non-concatemer (FLNC) reads were then subjected to de novo clustering with *isoseq cluster2*. This reference-free step groups reads originating from the same isoform to generate a highly accurate polished consensus sequence. These polished isoforms were subsequently aligned to the hg38 human reference genome (GENCODE v39) using *isoseq pbmm2*, an aligner optimized for long-read data. Finally, to create a non-redundant transcript set, *isoseq collapse* was used to merge isoforms that were splice-compatible, meaning they shared identical splice junctions but differed only in their 5’ or 3’ ends, which often reflects natural transcriptional start site variability.

### Classification and Filtering

With a set of unique isoforms defined, we performed structural annotation and rigorous quality control using the *isoseq pigeon* toolkit. The *pigeon classify* command categorized each isoform by comparing its splice junctions against the reference annotation, using the established SQANTI classification system (Pardo-Palacios et al. 2024). This process assigned labels such as Full Splice Match (FSM) for isoforms identical to known transcripts, Incomplete Splice Match (ISM) for partial matches, Novel in Catalog (NIC) for new combinations of known splice sites, and Novel Not in Catalog (NNC) for isoforms containing at least one previously unannotated splice site. To ensure the final dataset’s integrity, pigeon filter was applied to remove common long-read sequencing artifacts. This filtering specifically targeted isoforms resulting from intra-priming, where the oligo(dT) primer mistakenly anneals to an internal A-rich sequence instead of the true poly(A) tail, and reverse transcriptase (RT) template switching, which creates chimeric cDNAs.

The process also removed isoforms with poor read support for non-canonical splice junctions, yielding a final, high-confidence catalog of transcripts.

### Differential Expression and Functional Analysis

We performed differential expression analysis at both the gene and transcript levels using the edgeR v4.6 package in R (Chen et al. 2025). The analysis began with a read count matrix generated by the FLAIR v2.2 pipeline (Tang et al. 2020), which quantifies isoform expression from full-length transcript sequencing data. We derived gene-level counts by aggregating the counts of all constituent isoforms for each gene. To enhance statistical power and remove noise from lowly expressed features, we first filtered the data, retaining only those genes and transcripts that had at least two counts in a minimum of two samples. The core differential analysis was then conducted within the edgeR framework, which models the count data using a negative binomial distribution to appropriately account for the overdispersion typically seen in sequencing data. After normalizing for library size and composition using the Trimmed Mean of M-values (TMM) method, we fit a generalized linear model (GLM) to identify significant expression changes. Significance was determined based on a False Discovery Rate (FDR) cutoff of 0.05, which corrects for multiple hypothesis testing. To summarize the overall expression, change at the gene level while considering the contributions of different isoforms, we calculated a weighted mean log₂ fold change (LFC) for each gene. This was done by weighting the LFC of each individual isoform by its average normalized expression (baseMean), giving more influence on the more abundant isoforms. Finally, to understand the biological implications of these expression changes, we performed a functional enrichment analysis on the list of significant genes using the clusterProfiler package v4.0 (Wu et al. 2021).

### Differential Transcript Usage and Functional Consequence Analysis

This analysis was conducted using the IsoformSwitchAnalyzeR v2 R package(Vitting-Seerup and Sandelin 2019), a comprehensive toolkit for identifying and characterizing isoform switching events. We began by importing the isoform-level read counts, a GTF file defining transcript structures, and isoform nucleotide sequences generated by the FLAIR pipeline. These data sources were integrated into a switchAnalyzeRlist object, the central data structure for the analysis. A pre-filtering step was applied to remove genes with low expression, thereby increasing the statistical power to detect genuine switching events. The core statistical test for DTU was performed by leveraging the power of the DEXSeq package, which is designed to detect changes in feature usage relative to the parent gene’s total expression. This method, combined with limma’s empirical Bayes approach for variance moderation (Ritchie et al. 2015), with significance defined by FDR < 0.05.

Beyond statistical significance, we aimed to understand the functional consequences of these isoform switches. First, we predicted the Open Reading Frames (ORFs) for all isoforms to determine their protein-coding sequences. We then used a suite of specialized tools to predict functional attributes of the resulting proteins. This included assessing the coding potential of each transcript with CPAT v3.0.5(Wang et al. 2013), identifying conserved functional protein domains with Pfam v37.2 (Punta et al. 2012), and predicting signal peptides for secretion using SignalP v6.0(Della Marina et al. 2024). We also analyzed protein topology by predicting transmembrane helices with DeepTMHMM v1.0.24(“DeepTMHMM Predicts Alpha and Beta Transmembrane Proteins Using Deep Neural Networks - Abstract - Europe PMC,” n.d.), identified intrinsically disordered regions with IUPred2A (Mészáros et al. 2018), and predicted the final sub-cellular localization of the proteins with DeepLoc2 v2.1 (Thumuluri et al. 2022). Finally, we used the extractTopSwitches function to extract the most biologically relevant events. This function filters for switches that were not only statistically significant (FDR < 0.05) but also showed a substantial change in proportional expression (an isoform fraction difference > 0.1) and resulted in a predicted functional change.

### DDX5 Perturbation

#### gRNA design and cloning for DDX5

A plasmid containing the KRAB domain fused to dCas9 and the vector to clone sgRNA were obtained from Addgene (RRID: Addgene_99372 and Addgene_99376) (PMID:28757163). The plasmid #99376 contains a dTomato fluorescent protein. DDX5 sgRNA was custom designed by using the CHOPCHOP algorithm design tool. Forward and reverse guides of each target were used to generate duplexes and ligated into the *BsmBI* vector (Xing et al., 2014). The sgRNA cloned in the vector #99376 was confirmed by Sanger sequencing.

*lenti-EF1a-dCas9-KRAB-Puro was a gift from Kristen Brennand (Addgene plasmid #99372; http://n2t.net/addgene:99372; RRID: Addgene_99372)

### *lentiGuide-Hygro-dTomato was a gift from Kristen Brennand (Addgene plasmid #99376: http://n2t.net/addgene:99376; RRID: Addgene _99376)

### Overexpression of splicing genes

#### Cloning of NOVA1 and PTBP2 gene into TetO MCS

cDNA using SuperScript IV First Strand Synthesis System (Cat # 18091050) was synthesized using 1ug of total RNA. 100ng of the synthesized cDNA was used to set up a PCR using primer sequences that include:

For NOVA1 gene: NOVA1_XbaI_Kozak_Fw (5’ GAATCTAGAGCCACCATGATGGCGGCAGCTCCCAT 3’),

NOVA1_NheI_Rv (5’ AGAGCTAGCTCAACCCACTTTCTGAGGAT 3’). PCR was performed using Q5 Hot Start DNA <u>polymerase</u> (NEB M0493S) in 50uL reaction. Reaction parameters include: 10 cycles 64°C annealing, 60sec extension, 25 cycles 72°C annealing, 60sec extension, and primer concentration of 500nM each.

For PTBP2 gene: PTBP2_XbaI_Kozak_Fw (5’ GAATCTAGAGCCACCATGGACGGAATCGTCACTGA 3’),

PTBP2_NheI_Rv (5’ AGAGCTAGCTTAAATTGTTGACTTGGAGA 3’). PCR was performed using Q5 Hot Start DNA <u>polymerase</u> (NEB M0493S) in 50uL reaction. Reaction parameters include: 10 cycles 57°C annealing, 60sec extension, 25 cycles 68°C annealing, 60sec extension, and primer concentration of 500nM each.

PCR amplicons were excised from agarose gels and purified using the QIAquick Gel Extraction Kit (Cat. #28704). The purified products were cloned into the Zero Blunt TOPO PCR Cloning vector (Cat. #450245) according to the manufacturer’s protocol. Plasmids were isolated from selected TOPO clones by miniprep to obtain inserts for subsequent subcloning.

For restriction-based cloning, 2 µg of the TOPO plasmid containing the target gene and 1 µg of the FUW-tetO-MCS backbone (RRID: Addgene_84008; PMID: 27641305) were digested with XbaI and NheI. Digested products were purified and ligated at a 1:3 (vector: insert) molar ratio using T4 DNA ligase (NEB M0202S). Positive clones were screened by diagnostic restriction digestion, and insert identity was verified by plasmid sequencing (Plasmidsaurus).

#### Lentiviral preparation and infection

To prepare lentivirus, HEK293T cells (ATCC, cat #: CRL-11268) (RRID: CVCL_1926) at 60% confluency were transfected with 3rd generation packaging vectors (REV, RRE, and pMD 2.G) (RRID: Addgene_12253, Addgene_12251 and Addgene_12259), dCas9-KRAB and DDX5 gRNAs plasmids using the calcium phosphate method. 24 hours after transfection, the media was replaced. Virus was harvested 24 hrs after the media change (48 hrs after transfection). After filtering through 0.45μm low protein-binding syringe filters (SLHVM33RS Millipore), supernatant containing the viral particles was stored at −80℃. Cells were infected and treated with Puromycin 1μg/mL and Hygromycin 200 μg /mL concentration for 48 hrs. The antibiotics were added 48 hours after infection.

### Immunocytochemistry

Cells were first fixed using 4% paraformaldehyde (15710 Electron Microscopy Science) in PBS. The fixed samples were incubated with 0.5% Triton X-100 (T8787 Sigma-Aldrich) in PBS for 10 minutes. 5% FBS (100-106 Gemini Bio) in PBS was used for blocking for 1 hour at RT. Primary antibodies against Nanog (Abcam Cat# ab21624, RRID: AB_446437) for iPSCs marker and CNN (Sigma-Aldrich Cat# C2687, RRID: AB_476840) and TAGLN (Abcam Cat# ab14106, RRID: AB_443021) for VSMCs marker diluted in blocking buffer were incubated for approximately 2 hours at room temperature or overnight at 4°C. After three PBS washes, the secondary antibodies Donkey anti-mouse 488 (Alexa-fluor A32766 Invitrogen, RRID: AB_2762823), Donkey anti-rabbit 555 (Alexa-fluor A32794 Invitrogen, RRID: AB2762834), and DAPI (MBD0015 Sigma‒Aldrich) in blocking buffer were added for 1 hour at RT. Three additional PBS washes were performed, and the cells were imaged.

### RNA extraction, cDNA synthesis, and qPCR

The cell pellets were dissociated using TRIzol (15596018 Invitrogen). RNA was then extracted using a Zymo Direct-zol RNA Miniprep kit (R2052) according to the manufacturer’s protocol. iScript Reverse Transcription Reaction Mix (1708891, Bio-Rad) was used for cDNA synthesis. For qPCR, iTaq Universal SYBR Green Supermix (1725121 Bio-Rad) and SsoAdvanced Universal SYBR Green Supermix (17252721 Bio-Rad) was used on a Bio-Rad CFX OPUS 384 Touch Real-Time PCR machine. GraphPad Prism (RRID: SCR_002798) was used to perform statistical analysis using unpaired t-tests or 1-way ANOVA with Bonferroni correction. A complete list of primers used for qPCR is provided below.

### Statistical analysis

Details of statistical analyses are given in the Figure Legends. These include the number of samples and the number of independent experiments. Data were analyzed using unpaired t-test or one-way ANOVA with Bonferroni multiple comparison post hoc test as specified in the corresponding figure legends.

## Primer Table

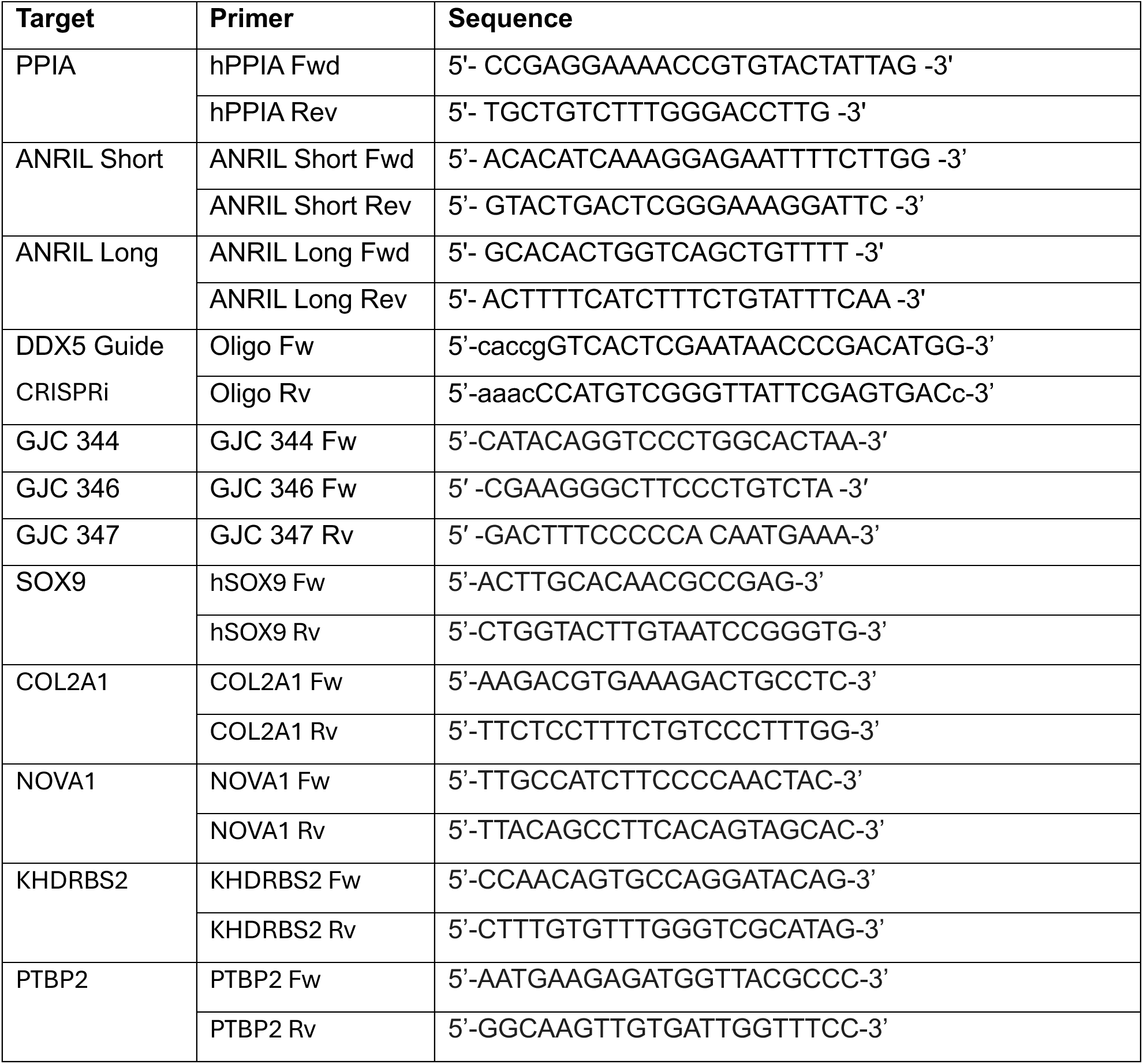
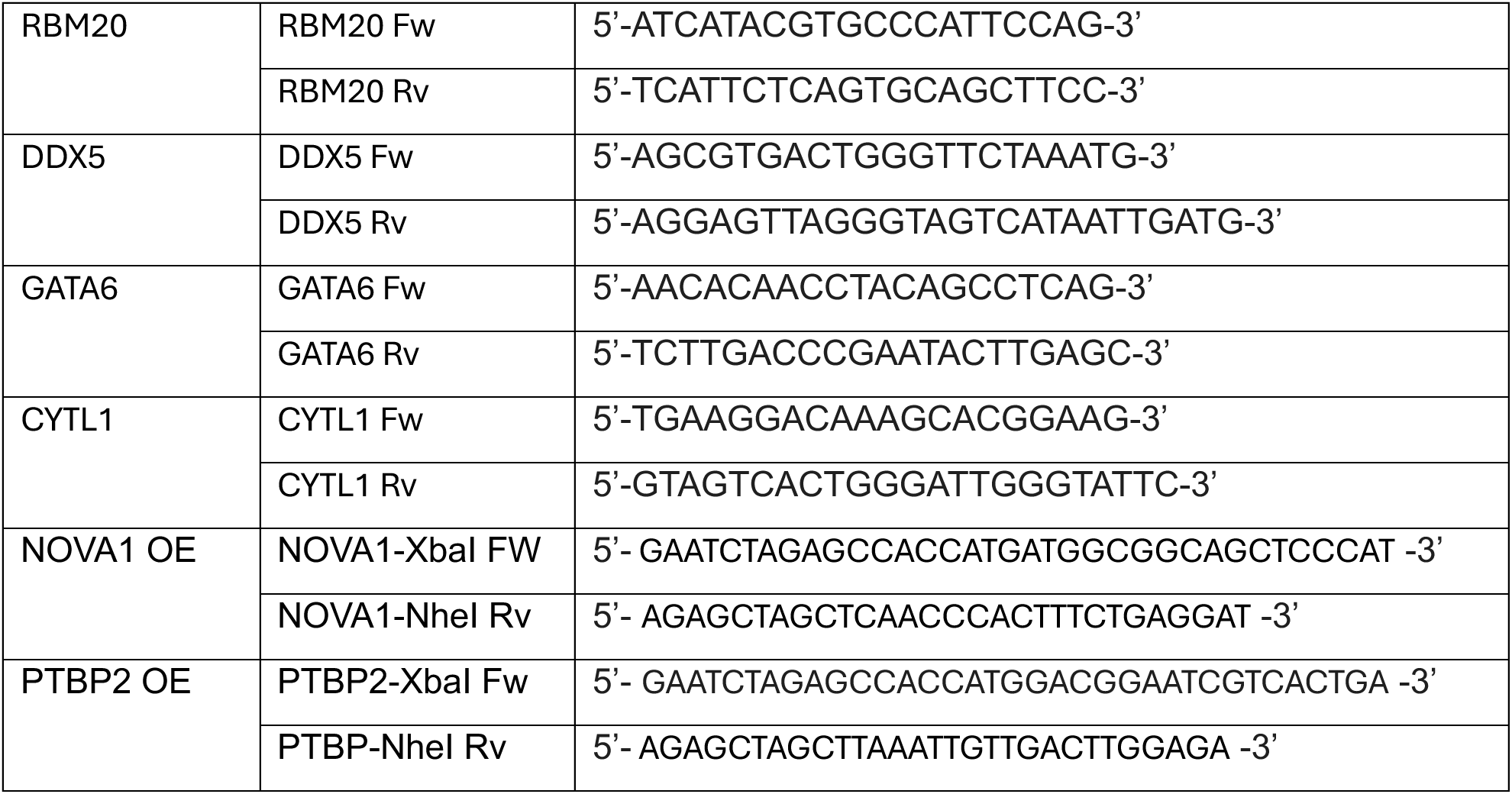

## Acknowledgments

The authors utilized the University of Wisconsin – Madison Biotechnology Center Gene Expression Center Core Facility (Research Resource Identifier – RRID:SCR_017757).

## Author contribution

Conceptualization, S.S., H.Y. and V.L.S. Data curation, S.S. and H. Y. Formal analysis and Methodology, S.S., H.Y., E. S. and K.M. Manuscript writing and data visualization S.S, H.Y and V.L.S.

## Source of funding

The authors acknowledge the following funding: S.S. was supported by the UW Stem Cell and Regenerative Medicine Center Graduate training award. E.S. was supported by the NIH T32 Genetics Training grant (T32GM007133) and Advanced Opportunity Fellowship through SciMed Graduate Research Scholars, and the CVRC T32 at the University of Wisconsin Madison. This work was supported by the National Institute of Health grants R35GM155127 (VLS) and the Wisconsin Partnership Program New Investigator Award (VLS). V.L.S. also received funding for this study from the University of Wisconsin Madison School of Medicine and Public Health.

## Disclosure

Nothing to disclose.

---

**Figure S1.**
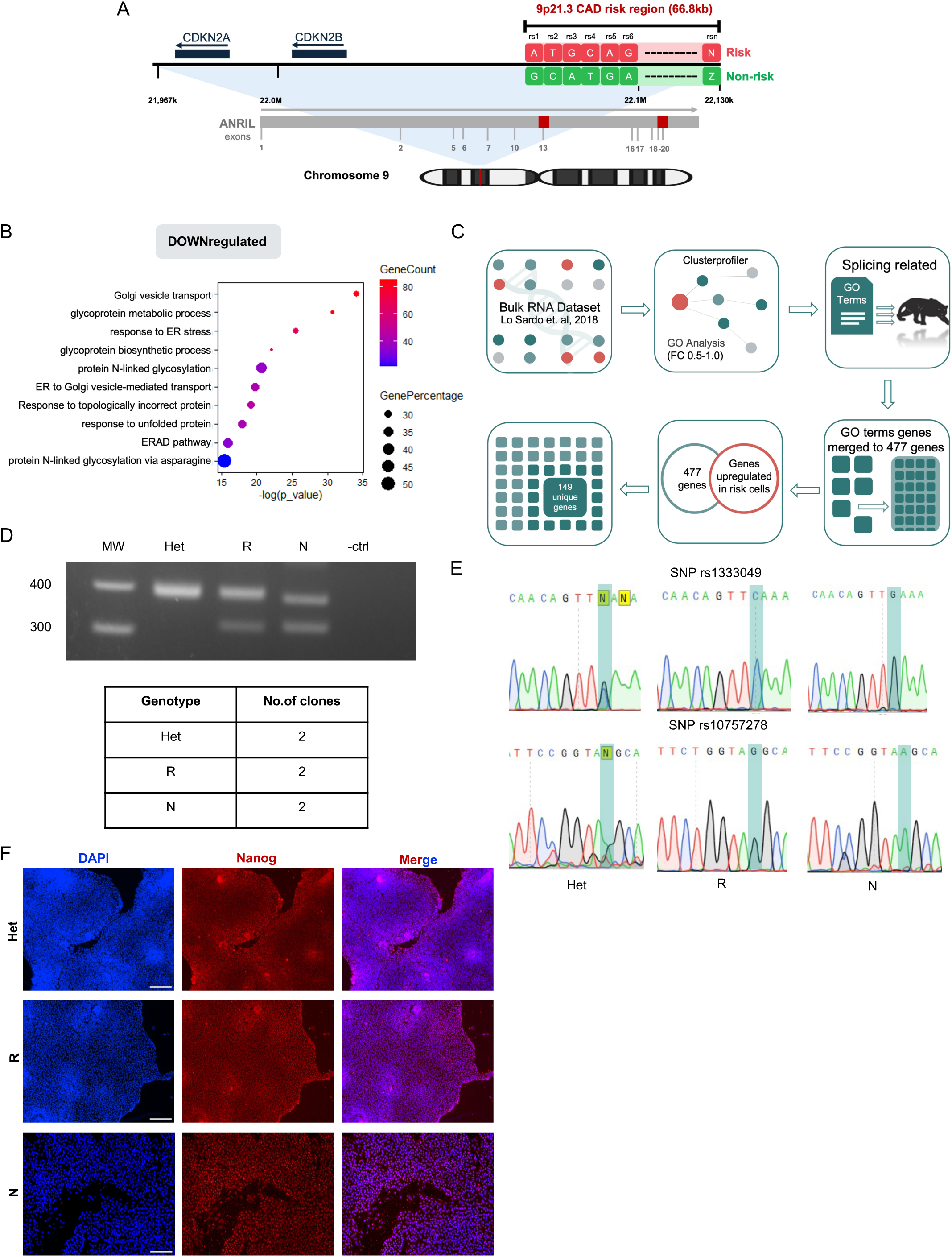
Evaluation of splicing genes alteration and validation of the new isogenic iPSC collection. **(A)** Schematic of the 9p21.3 coronary artery disease locus. The cartoon shows the human chromosome 9 and the 66.8 Kb CAD risk region. The CAD locus is flanked by the CDKN2A and B genes and overlaps with the ANRIL gene. Red square in the ANRIL gene indicate the two sites for termination of transcription in exon 13 and 20. Within the CAD region a simplified schematic of the different pattern of SNPs (rs) at the region resulting in two possible alleles (risk and non-risk). **(B)** The top 10 gene ontology terms downregulated in the differentially expressed genes in RR VSMCs between log_2_ fold change 0.1- 1.0. **(C)** Schematic of workflow to prioritize genes related to splicing**. (D)** R, N, and Het iPSCs were genotyped for the deletion by PCR as we previously described (Lo Sardo et al 2018). 2 clones per genotype were generated and used for experiments. **(E)** The genotype of the newly generated iPSCs was confirmed by Sanger sequencing for two SNPs at the 9p21.3 CAD locus (rs1333049, rs10757278). **(F)** Immunocytochemistry for the pluripotency marker Nanog for Het, R, and N iPSCs. Scale is 100 μm.

**Figure S2.**
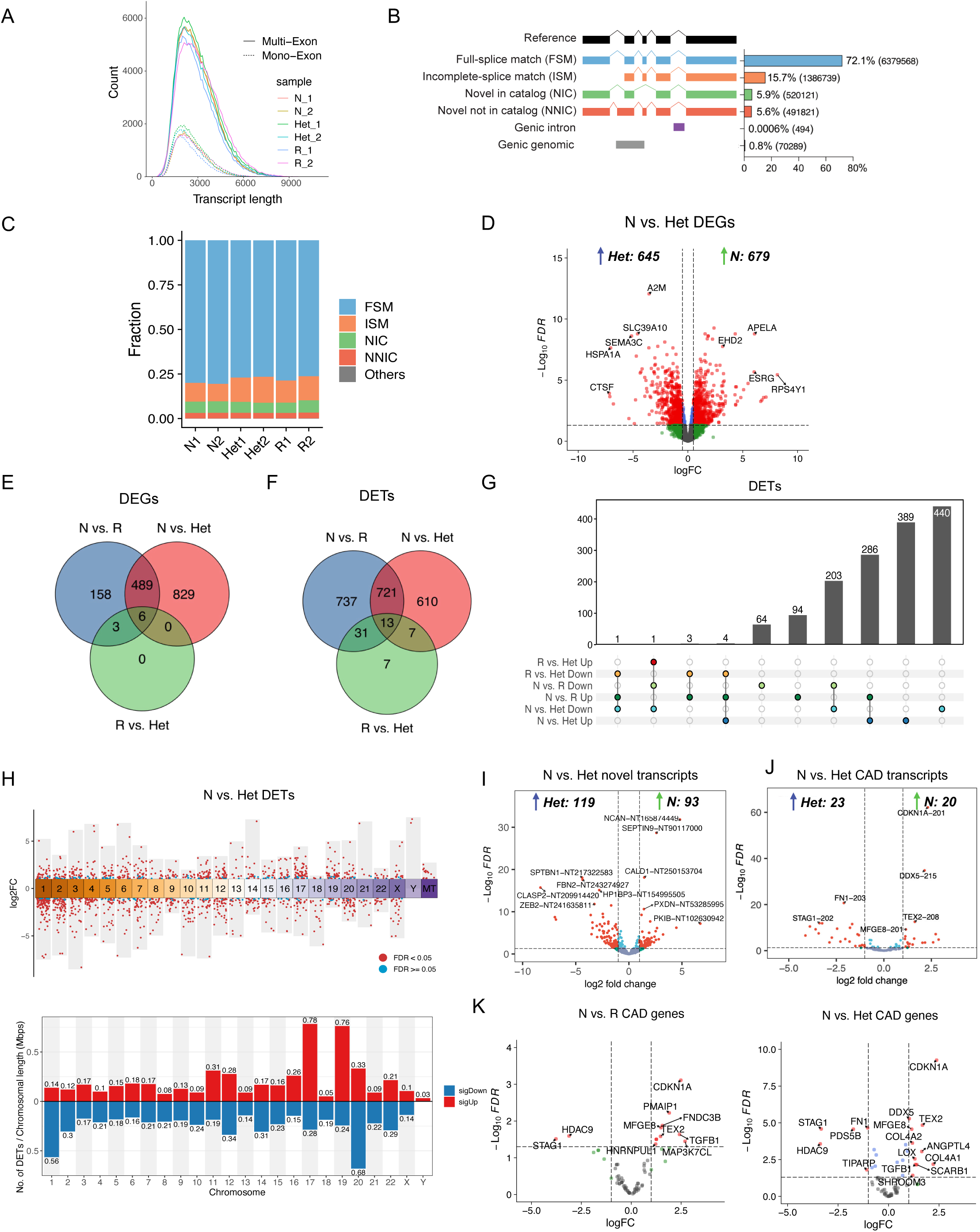
Pac-Bio long read sequencing of VSMCs with different haplotypes at the 9p21.3 CAD locus. **(A)** Distribution of transcript lengths across samples, showing multi-exon (solid lines) and exon (dashed lines) transcripts. **(B)** Classification of transcripts based on splice junction matching to known annotations, including full-splice match (FSM), incomplete-splice match (ISM), novel isoforms in catalog (NIC), and novel not in catalog (NNIC), along with genic and intergenic categories. Percentages represent the proportion of each class. **(C)** Fraction of isoform types in each sample, demonstrating the relative abundance of FSM, ISM, NIC, NNIC, and other categories across all biological replicates. **(D)** Volcano plots illustrate genome-wide differential gene expression between N vs. Het VSMCs. logFC: log2 fold change. **(E)** Venn diagram representing the overlap of DEGs (FDR < 0.05 and log_2_FC > 1 or <-1) across the three genotypes. **(F)** Venn diagram representing the overlap of DETs (FDR < 0.05 and log_2_FC > 1 or <-1) across the three genotypes. **(G)** Upset plot represents the intersection structure of DETs. Bars represent the size of DET subsets, while the connected dots below each bar indicate which comparisons contribute to each subset. **(H)** Chromosomal distribution of DETs for log_2_ fold changes across the genome for N vs. Het haplotypes. Each dot represents a transcript positioned along chromosomes 1–22, X, and Y, as well as mitochondrial DNA. Bar plots showing the number of DETs per chromosome normalized to chromosome length, for N vs Het. Red bars indicate upregulated DETs, blue bars indicate downregulated DETs. **(I)** Volcano plots showing differential expression of novel transcripts (NIC and NNIC), for N vs Het VSMCs. **(J)** Volcano plots showing differential expression of CAD-associated transcripts, for N vs Het VSMCs. **(K)** Volcano plots showing differential expression of CAD-associated genes, for N vs R VSMCs and N vs Het.

**Figure S3.**
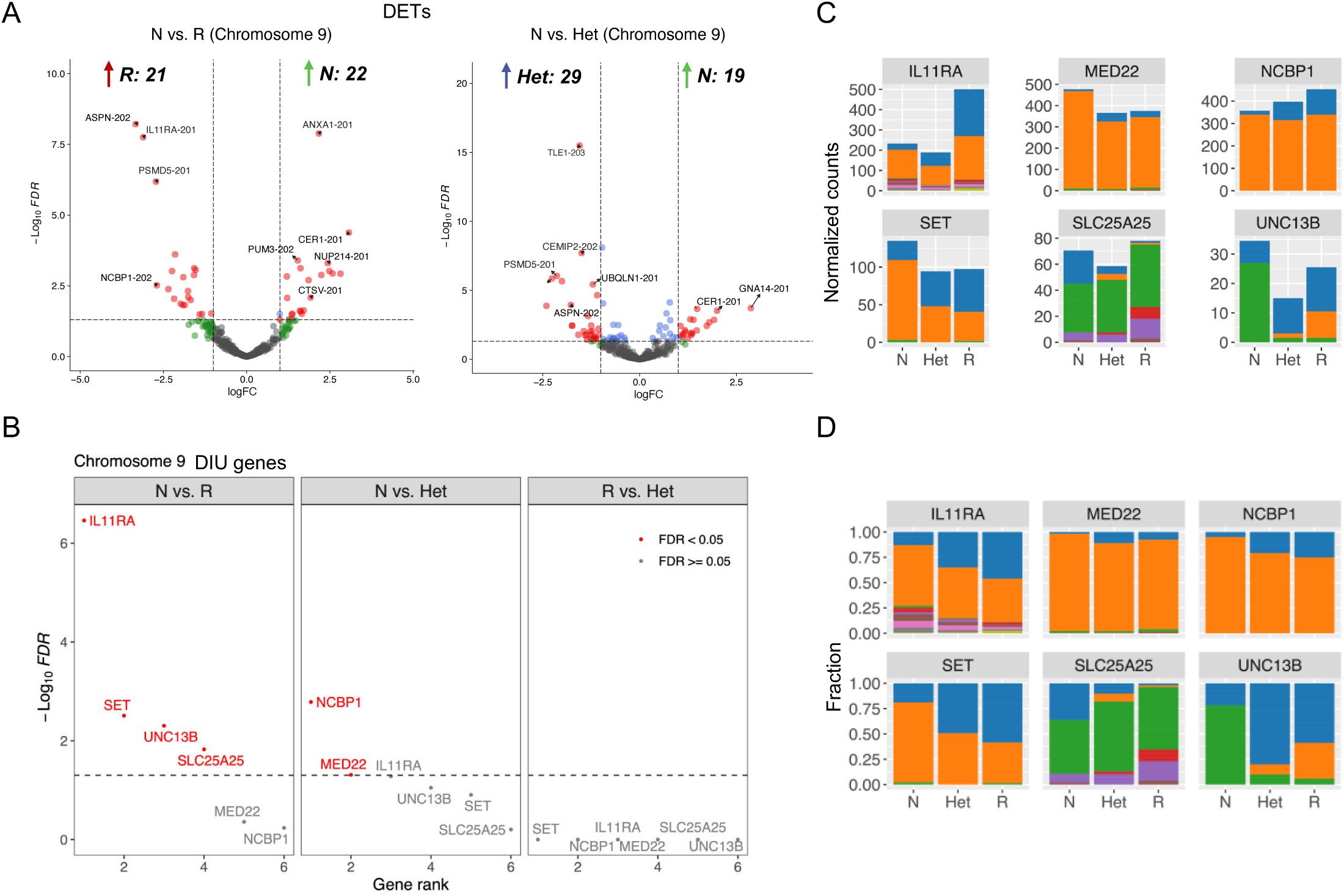
Differential transcripts analysis at chromosome 9. **(A)** Volcano plots illustrate the differential expression of transcripts in chromosome 9. **(B)** Genes showing differential isoform usage (DIU) are ranked based on the -log10 FDR of DIU. **(C-D)** The DIU of significant isoform switch genes is presented using normalized counts **(C)** and fraction **(D)**. Each color represents a distinct transcript isoform.

**Figure S4.**
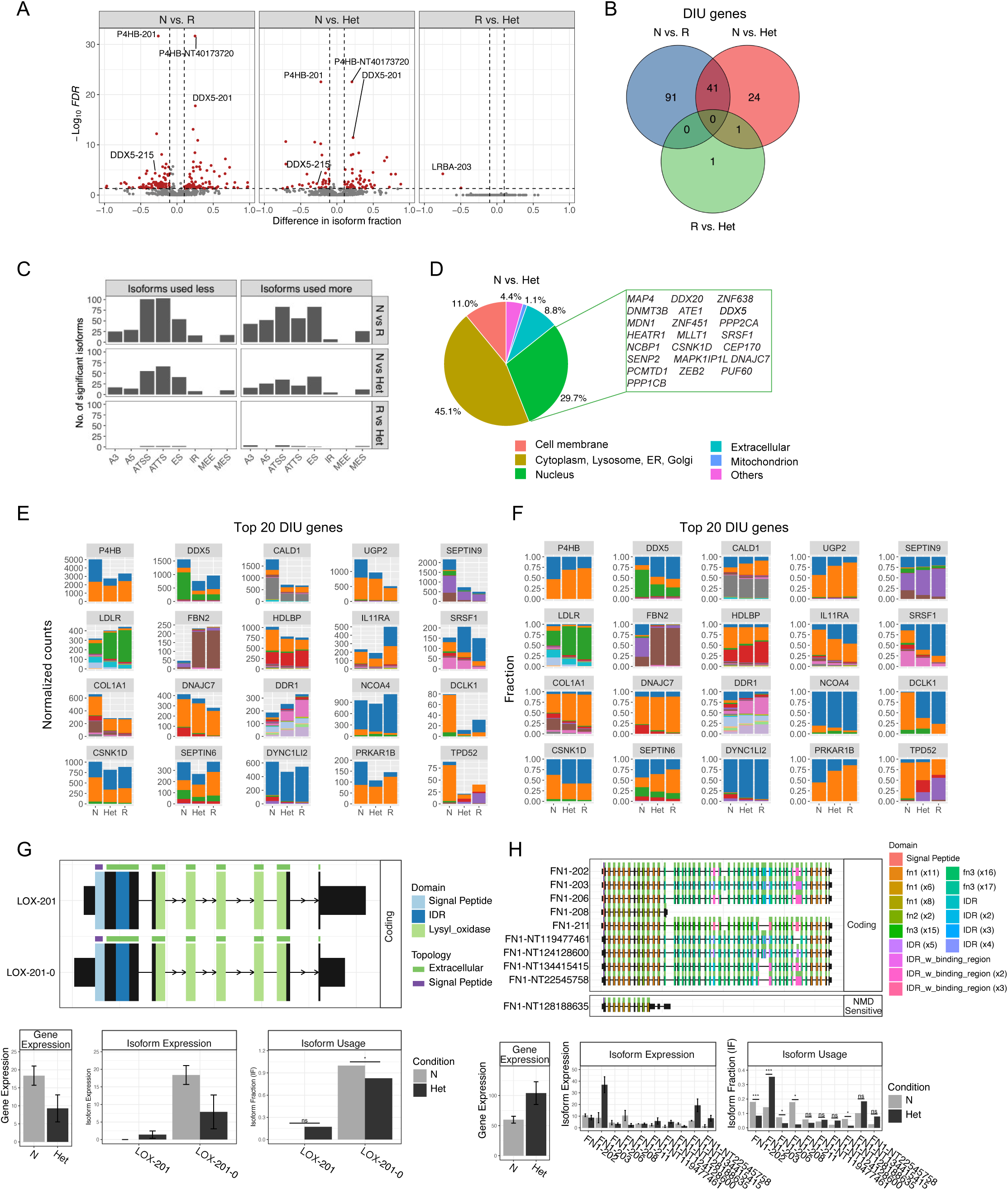
Differential transcript usage driven by different haplotypes at 9p21.3. **(A)** Volcano plots showing differential isoform usage between Non-risk (N), Risk (R), and Heterozygous (Het) groups. Red dots indicate isoforms with significant changes (FDR < 0.05). **(B)** Venn diagram representing the overlap of DTUs across the three genotypes. **(C)** Bar charts showing the distribution of alternative splicing event types among isoforms used less **(left)** and more **(right)** in each genotype comparison. **(D)** Pie charts representing the distribution of cellular localization of proteins encoded by genes undergoing significant isoform switching between N vs. Het haplotypes. **(E-F)** DIU of the top 20 genes by normalized counts **(E)** and fractions **(F)** between genotypes. Each color represents a distinct transcript isoform. **(G)** Schematic representation showing distinct exon composition and domain architecture of LOX protein isoforms. Bar graphs demonstrate gene-level expression, isoform-specific expression, and isoform usage of LOX between N vs. Het. **(F)** Schematic representation showing distinct exon composition and domain architecture of FN1 protein isoforms. Bar graphs demonstrate gene-level expression, isoform-specific expression, and isoform usage of FN1 between N vs. Het.

**Figure S5.**
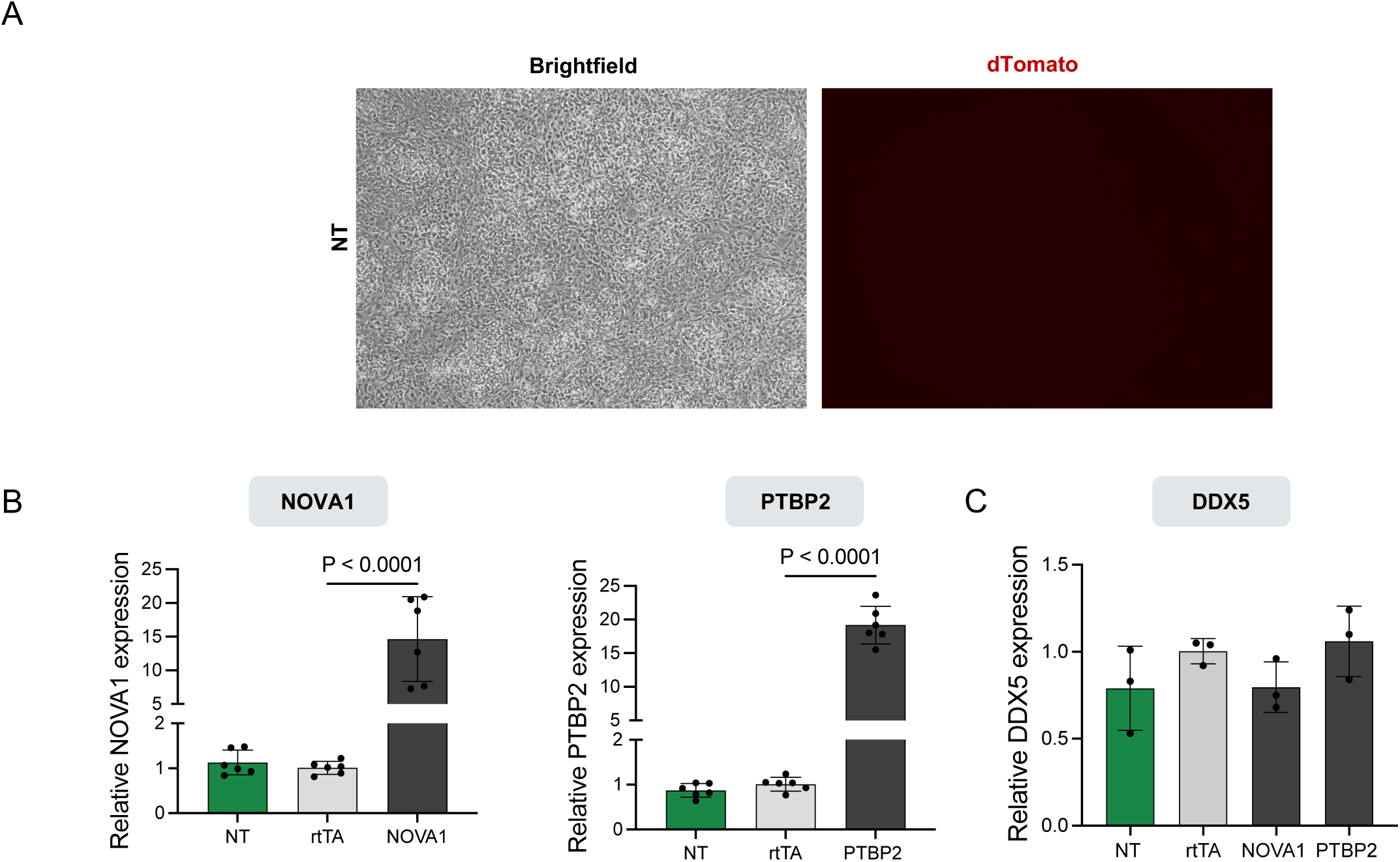
DDX5 expression is independent of splicing genes. **(A)** Representative brightfield and fluorescence images of VSMCs from the DDX5i experiments with no viral infection (NT) relative to Figure 4D. Scale is 200 μm. **(B)** qPCR quantification of overexpression of splicing genes NOVA1 and PTBP2 **(C)** qPCR quantification of DDX5 in cells overexpressing the splicing genes NOVA1 and PTBP2. P-value calculated with One-way ANOVA with Bonferroni correction.

## Notes

### Competing Interest Statement

The authors have declared no competing interest.

### Summary of Updates

The text of the manuscript has been revised to convey a more clear message of the study and enhance the findings for a broader impact.

